# Biallelic *TANGO1* mutations cause a novel syndromal disease due to hampered cellular collagen secretion

**DOI:** 10.1101/750349

**Authors:** Caroline Lekszas, Ombretta Foresti, Ishier Raote, Daniel Liedtke, Eva-Maria König, Indrajit Nanda, Barbara Vona, Peter De Coster, Rita Cauwels, Vivek Malhotra, Thomas Haaf

**Author notes:** These authors contributed equally to this work (Caroline Lekszas, Ombretta Foresti, Ishier Raote). These authors also contributed equally to this work (Vivek Malhotra, Thomas Haaf). Caroline Lekszas, Ombretta Foresti, Ishier Raote, Daniel Liedtke, Eva-Maria König, Indrajit Nanda, Barbara Vona, Peter De Coster, Rita Cauwels, Vivek Malhotra, Thomas Haaf. Corresponding author: Thomas Haaf.

## Abstract

The transport and Golgi organization 1 (TANGO1) family proteins have been shown to play pivotal roles in the secretory pathway. Full length TANGO1 is a transmembrane protein localised at endoplasmic reticulum exit sites (ERES), where it binds bulky cargo within the ER lumen and recruits membranes from the ER Golgi intermediate compartment (ERGIC) to create an exit route for their export. *Tango1* knockout mice display a global collagen secretion defect and perinatal lethality. Here we report the first TANGO1-associated syndrome in humans, which mainly manifests in a collagenopathy. A synonymous substitution that results in exon 8 skipping in most mRNA molecules, ultimately leading to a truncated TANGO1 protein was identified as the disease-causing mutation. The four homozygously affected sons of a consanguineous family display severe dentinogenesis imperfecta, short stature, various skeletal abnormalities, insulin-dependent diabetes mellitus, sensorineural hearing loss, and mild intellectual disability. Functional studies in HeLa and U2OS cells revealed that the corresponding truncated TANGO1 protein is dispersed in the ER and its expression in cells with intact endogenous TANGO1 impairs cellular collagen I secretion.

## Introduction

Collagens are the most abundantly secreted molecules in mammals and needed throughout the whole body for bone mineralization, skin and tissue assembly. Within the endoplasmic reticulum (ER) lumen, newly synthesized procollagens assemble into rigid, rod-like triple helices that are too large for export by the conventional coat protein complex II (COPII)-coated vesicle size of 60-90 nm (*Malhotra and Erlmann, 2015*; *Miller and Schekman, 2013*). Over the past years, the ER exit site (ERES) located, transport and Golgi organization 1 protein (TANGO1) has been identified as the key player in the export of hard to fold and bulky cargoes like the collagens (*Bard et al., 2006*; *Malhotra and Erlmann, 2011, 2015*; *Saito et al., 2009*; *Santos et al., 2016*; *Raote et al., 2017*, 2018; *Raote and Malhotra, 2019*; *Wilson et al., 2011*). *TANGO1* is conserved throughout most metazoans and ubiquitously expressed in humans. It comprises 8,142 bp located at chromosome 1q41 and encodes two distinct isoforms, full length TANGO1 and TANGO1-short. Full length TANGO1 consists of 1,907 amino acids (aa) and contains an N-terminal signal sequence followed by an Src-homology 3 (SH3)-like domain and a coiled-coil domain in the lumenal portion, as well as two additional coiled-coil domains (CC1 and CC2) and a proline-rich domain (PRD) in the cytoplasmic portion (*Figure 1A*). TANGO1-short is composed of 785 aa and lacks the lumenal portion contained in TANGO1 (*Saito et al., 2009*). Together with cTAGE5 encoded by the TANGO1-like protein gene (*TALI*), TANGO1 and TANGO1-short form stable complexes at ERES, to jointly fulfill their roles in the secretion of bulky cargoes such as procollagens, pre-chylomicrons, and large pre-very low-density-lipoproteins (*Bosserhoff et al., 2003*; *Maeda et al., 2016*; *Malhotra and Erlmann, 2011, 2015*; *Malhotra et al., 2015*; *Saito et al., 2009*, 2011; *Santos et al., 2016*; *Wilson et al., 2011*).

**Figure 1.**
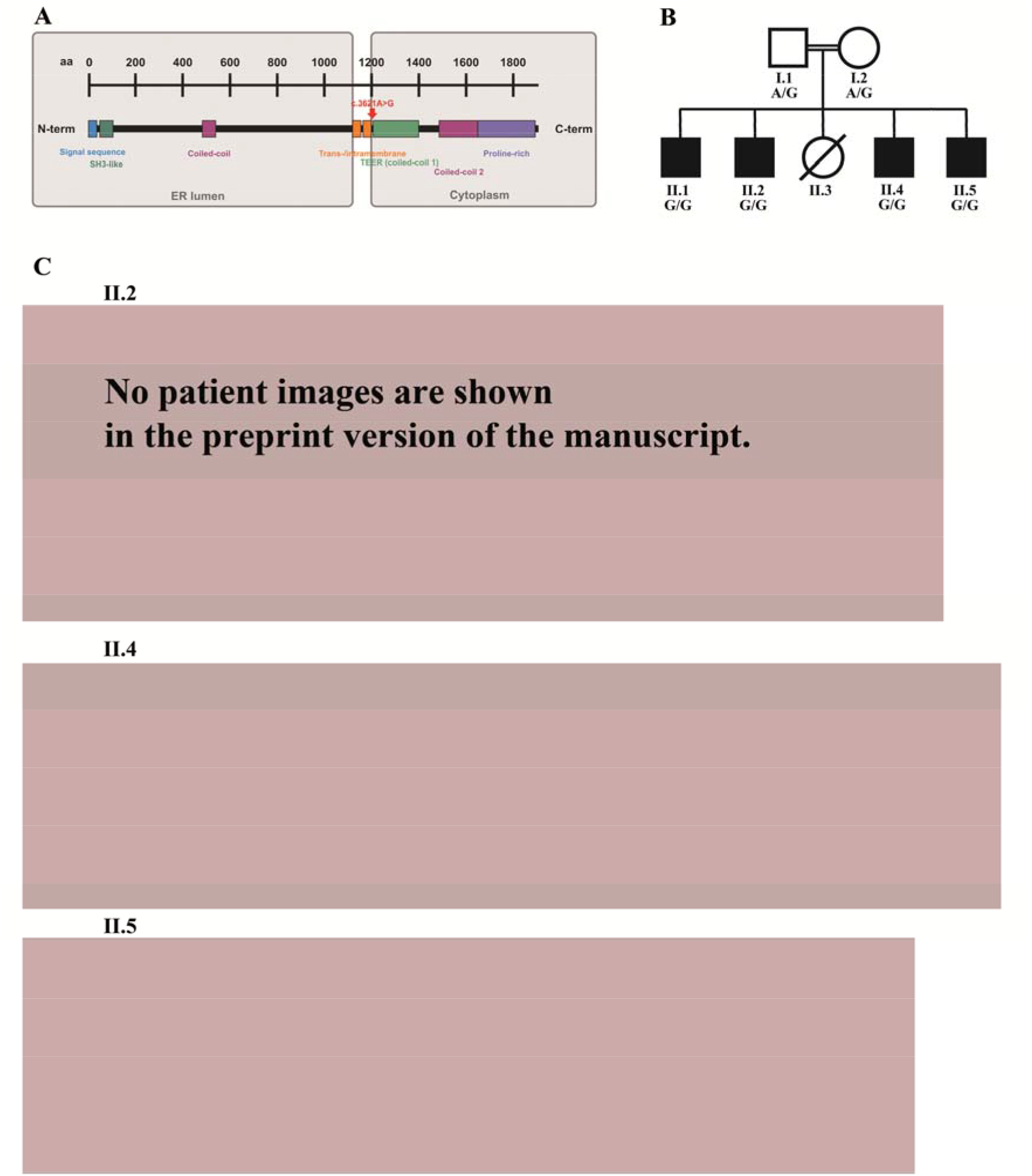
A novel collagenophathy caused by biallelic *TANGO1* mutations in a consanguineous family. (**A**) Structure of TANGO1 protein. The lumenal portion contains an N-terminal signal sequence followed by an SH3-like domain required for cargo binding, as well as a coiled-coil domain. A trans- and intramembrane domain anchors TANGO1 within the ER membrane. The cytoplasmic portion consists of two coiled-coil domains (CC1, also named TEER, and CC2) and a proline-rich domain at the C-terminus. The identified mutation affects residue 1207 (p.(Arg1207=)) between the intramembrane and the CC1 domain at the beginning of the cytoplasmic portion. (**B**) Pedigree of the studied family. The parents (I.1 and I.2) are first cousins. The four affected sons (II.1, II.2, II.4, and II.5) share a homozygous *TANGO1* (c.3621A>G) variant. The healthy child II.3 died in a household accident at the age of 16. (**C**) Phenotypic appearance of the affected brothers II.2, II.4, and II.5. Note the brachydactyly of hands and feet, clinodactyly of the fifth finger, high nasal bridge, dentinogenesis imperfecta (including an opalescent tooth discoloration with severe attrition affecting the primary and permanent dentition, as well as juvenile periodontitis, bulbous crowns, long and tapered roots, and obliteration of the pulp chamber and canals in the permanent dentition), the skin lesions due to pruritus in all affected children depicted; the scoliosis in II.4 and II.5; and the retrognathia in II.5.

At ERES TANGO1 assembles into rings that enclose COPII coats and create a sub-compartment dedicated to sorting, packing and exporting collagens (*Raote et al., 2017*, 2018; *Raote and Malhotra, 2019*). TANGO1’s SH3-like domain binds collagens via the collagen-specific chaperone HSP47 (heat shock protein 47) in the ER lumen (*Ishikawa et al., 2016*). This binding of TANGO1 to HSP47-Collagen is proposed to trigger binding of its PRD to Sec23 in the cytoplasm. TANGO1’s CC1 domain, that contains a subdomain named TEER (tether of ER Golgi intermediate compartment at ER), recruits ERGIC-53 membranes, which fuse with the nascent vesicle bud initiated by COPII inner coats (Sec23/Sec24) to grow the collagen filled container into an export conduit (*Santos et al., 2015; Raote et al., 2018; Raote and Malhotra, 2019*). Subsequent to collagen packing into this conduit, TANGO1 dissociates from HSP47 and collagen. TANGO1 is retained at ERES while collagens move forward in the anterograde direction (*Raote and Malhotra, 2019*).

The discovery of TANGO1 has made the process by which cells organize ERES and export collagen amenable to molecular analysis. We now describe the first human *TANGO1* mutation associated with a novel autosomal-recessive syndrome. These findings underscore the importance of TANGO1 in human (patho)physiology.

## Results

### Clinical description

Four brothers with a similar combination of congenital anomalies (two of whom have already been described by *Cauwels et al., 2005*) were referred for oral examination to the Centre for Special Care, Ghent University Hospital, at the ages of 7 (*Figure 1B*; II.1;*1988), 3 (II.2;*1990), 6 (II.4;*2006), and 4 (II.5;*2008) years, and were followed up until present. Their parents are of Turkish origin and first cousins. The sister (II.3) as well as both parents (I.1 and I.2) were phenotypically normal. All four brothers presented with severe dentinogenesis imperfecta in both primary and permanent dentitions, delayed eruption of the permanent teeth, growth retardation, proportionate short stature, clinodactyly of the fifth finger, brachydactyly, platyspondyly, primary obesity, insulin-dependent diabetes mellitus (<1 IU of insulin/kg/day), sensorineural hearing loss, and mild intellectual disability (ID). Additional facultative symptoms included scoliosis, retrognathia, mild retinopathy, osteopenia, early onset periodontitis with premature tooth loss, hydronephrosis, and microalbuminuria (*Figure 1C*; *Table 1*).

**Table 1.**
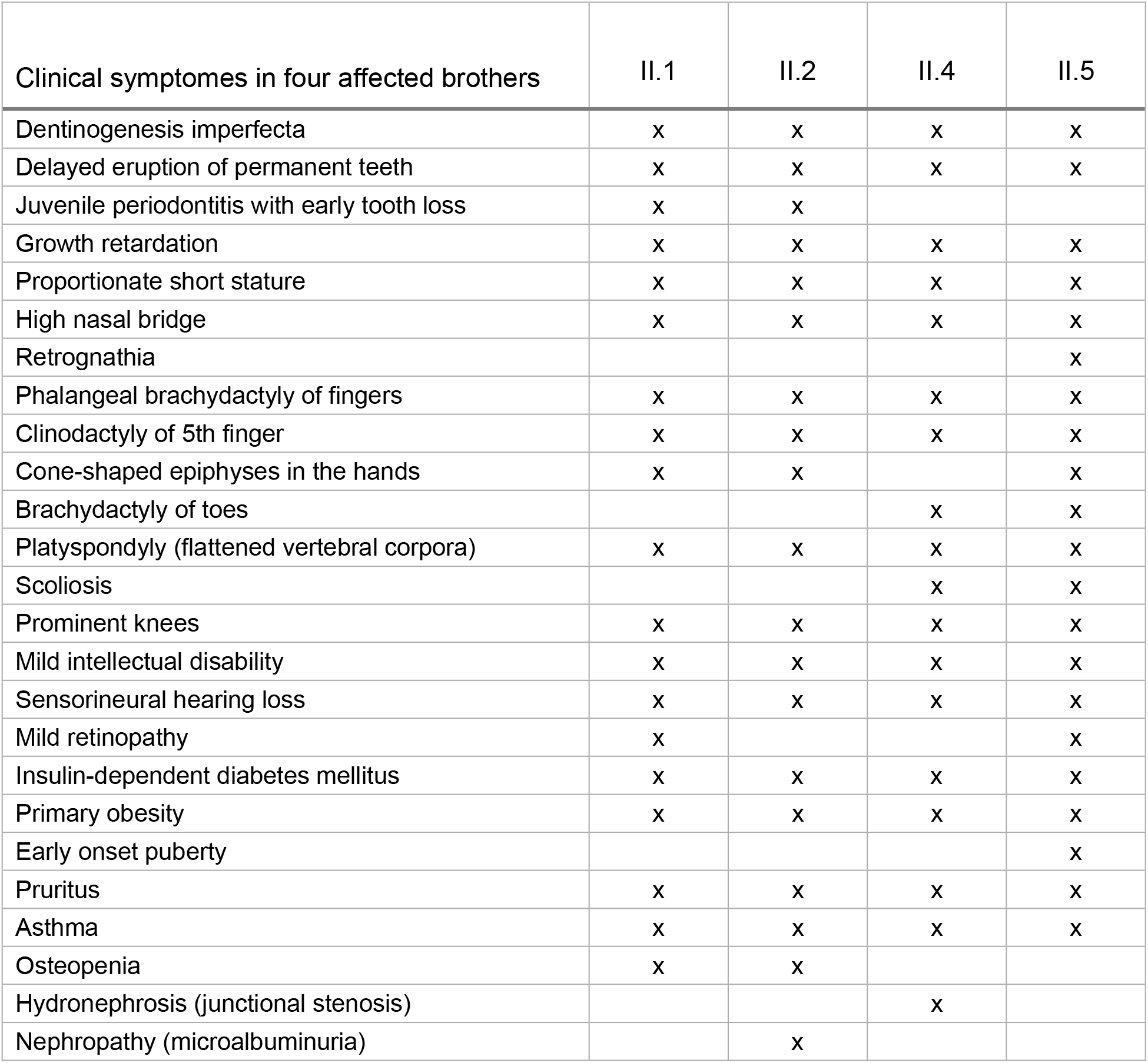

| **Table 1.**<br>Clinical symptoms in four affected brothers | II.1 | II.2 | II.4 | II.5 |
| --- | --- | --- | --- | --- |
| Dentinogenesis imperfecta | x | x | x | x |
| Delayed eruption of permanent teeth | x | x | x | x |
| Juvenile periodontitis with early tooth loss | x | x |  |  |
| Growth retardation | x | x | x | x |
| Proportionate short stature | x | x | x | x |
| High nasal bridge | x | x | x | x |
| Retrognathia |  |  |  | x |
| Phalangeal brachydactyly of fingers | x | x | x | x |
| Clinodactyly of 5th finger | x | x | x | x |
| Cone-shaped epiphyses in the hands | x | x |  | x |
| Brachydactyly of toes |  |  | x | x |
| Platyspondyly (flattened vertebral corpora) | x | x | x | x |
| Scoliosis |  |  | x | x |
| Prominent knees | x | x | x | x |
| Mild intellectual disability | x | x | x | x |
| Sensorineural hearing loss | x | x | x | x |
| Mild retinopathy | x |  |  | x |
| Insulin-dependent diabetes mellitus | x | x | x | x |
| Primary obesity | x | x | x | x |
| Early onset puberty |  |  |  | x |
| Pruritus | x | x | x | x |
| Asthma | x | x | x | x |
| Osteopenia | x | x |  |  |
| Hydronephrosis (junctional stenosis) |  |  | x |  |
| Nephropathy (microalbuminuria) |  | x |  |  |

### Whole exome sequencing (WES) revealed the disease-causing mutation in *TANGO1*

WES was performed in the four affected bothers and their parents. After filtering, 10 variants were found to be homozygous in all affected children and heterozygous in both parents. In-depth data analysis revealed a synonymous variant in exon 8 of *TANGO1* (NM_001324062.1: c.3621A>G) as the most likely disease-causing mutation. *TANGO1* is known to be crucial for the secretion of collagens, consistent with a collagenopathy in our patients. WES results were validated by Sanger sequencing (*Figure 2A*). This *TANGO1* mutation was not present in large population databases such as ExAC or gnomAD. Although it does not alter the amino acid at the respective position (p.(Arg1207=)), the A>G substitution was predicted by ESEfinder to disrupt an exon splice enhancer (ESE) motif recognized by the human SR protein SC35 (*Figure supplement 1A*). The mutation affects residue 1,207 between the intramembrane and the CC1 domain within the cytoplasmic portion of TANGO1 (*Figure 1A*).

**Figure 2.**
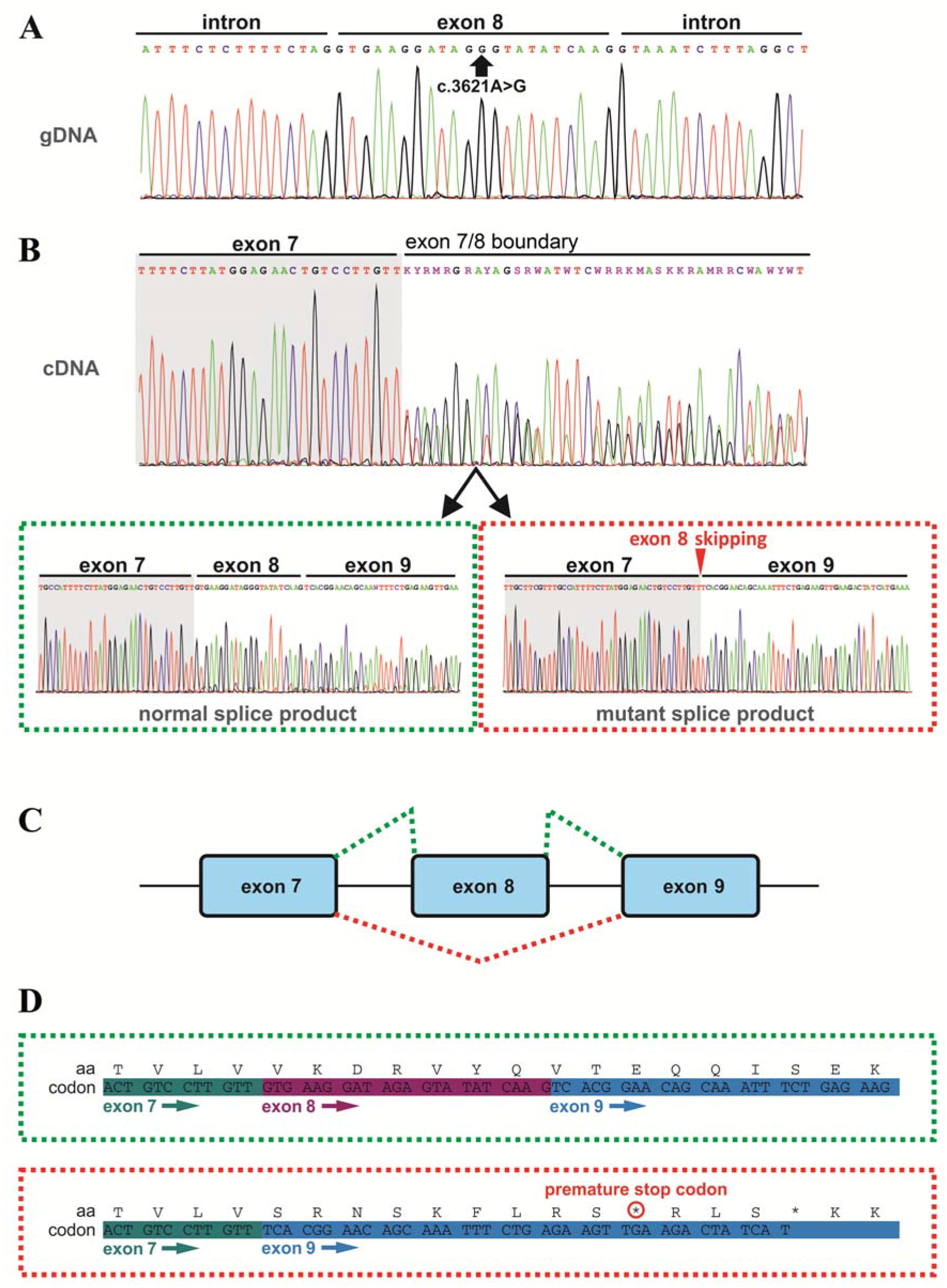
Effects of the *TANGO1* (c.3621A>G) mutation on pre-mRNA splicing. (**A**) The synonymous variant resides in exon 8 of *TANGO1* at genomic position 222,822,182 (GRCh37/hg19). It is predicted to disrupt an exon splice enhancer (ESE) motif recognized by the human SR protein SC35. (**B**) Electropherograms of the *TANGO1* cDNA sequence of one affected child. Note the splitting of the sequence starting at the exon 7/8 boundary. Sequencing of individual bands after gel electrophoretic separation revealed *TANGO1* wild-type cDNA and cDNA lacking exon 8 (c.3610_3631delins30). (**C**) Schematic representation of the alternatively used splice sites resulting in the normal *TANGO1* mRNA (green dotted lines) and in exon 8 skipping (red dotted lines). (**D**) Consequences of *TANGO1* exon 8 skipping on the reading frame and the amino acid level. Exclusion of exon 8 causes a premature stop codon.

Homozygosity mapping of two affected brothers (II.1 and II.2) identified a shared homozygous interval of ∼19 Mb on chromosome 1, spanning GRCh37/hg19 coordinates 214,413,099-233,429,284 (rs12736101-rs6656327), including *TANGO1* and 28 disease-causing OMIM genes (*Figure supplement 1B,C*). Apart from *TANGO1*, none of the shared homozygous intervals was endowed with a pathogenic mutation. In addition, no potentially disease-causing copy number variation (CNV) was detected by array comparative genomic hybridization (CGH).

### The *TANGO1* mutation leads to exon 8 skipping by disrupting an exon splice enhancer both *in vivo* and *in vitro*

In order to investigate possible effects of the identified *TANGO1* mutation on pre-mRNA splicing, blood samples of the whole family were used for RNA isolation and reverse transcription PCR. Subsequent gel electrophoresis and cDNA sequencing revealed two splice products in the homozygous sons and their heterozygous parents, one representing the full length transcript being more abundant in the parents and another one lacking the entire exon 8 being more abundant in the affected sons (*Figure 2B,C*; *Figure 3*). Exon 8 skipping during *TANGO1* pre-mRNA splicing causes a frameshift and a premature stop codon 27 bp downstream of exon 7 (c.3610_3631delinsTCACGGAACAGCAAATTTCTGAGAAGTTGA) (*Figure 2D*). The predicted truncated protein lacks almost the entire cytoplasmic portion including CC1/TEER, CC2, and PRD (*Figure 1A*).

**Figure 3.**
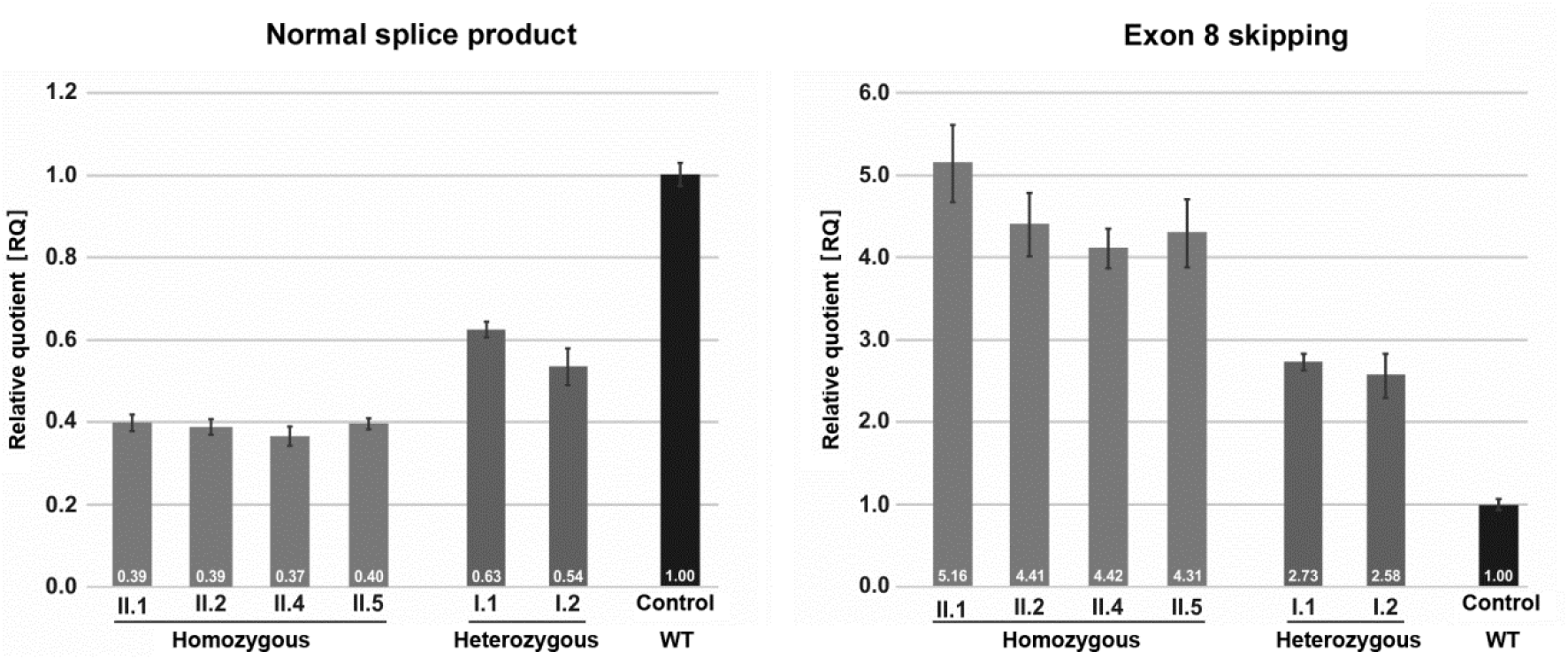
Quantification of *TANGO1* splice products in homozygous and heterozygous mutation carriers, compared to a control individual (without mutation). The right bar diagram shows the relative amounts of the normally spliced *TANGO1* cDNA and the left diagram of splice products lacking exon 8. A control cDNA sample was used for normalisation and relative comparison (RQ=1).

A minigene assay (*Figure supplement 2*) was performed to confirm that the identified *TANGO1* mutation is sufficient to induce exon 8 skipping. Transfection of cultured HEK293T cells with the Mnt vector carrying the mutated (c.3621A>G) *TANGO1* exon 8 and 250 bp flanking intronic sequences mainly produced vector-derived splice products lacking exon 8.

The observed splicing error could be due to disruption of an ESE motif recognized by the SR protein SC35 or to formation of a splice repressor motif recruiting the heterogeneous nuclear ribonucleoprotein A1 (hnRNP A1). To test this, cultured HeLa cells were transfected with either a wild-type or a mutated *TANGO1* vector (*Figure supplement 3*) and then treated with a customized antisense oligonucleotide (*vivo* morpholino) targeting the entire *TANGO1* exon 8. SR proteins are required for proper exon inclusion during splicing and their absence can lead to exon skipping. The observed effects of the morpholino treatment on *TANGO1* splicing support the idea that the c.3621A>G mutation interferes with SR protein binding.

### The homozygous *TANGO1* mutation results in exon 8 skipping in most splice products

Quantitative real-time (qRT) PCR on blood cDNA samples was performed to quantify the amount of mutant and normal *TANGO1* splice products, respectively, in homozygous and heterozygous mutation carriers compared to a normal control (*Figure 3*). The affected children consistently displayed the lowest amounts of normal splice product (mean RQ value: 0.39) and the highest amounts of the exon 8 skipped product (mean RQ value: 4.58). Both parents showed more normal splice product (mean RQ value: 0.59) than their children but still only approximately half of that of the control. In contrast to the control individual, both parents also displayed a considerable proportion of the exon 8 skipped splice product (mean RQ value: 2.66). Because of a processed transcript (ENST00000495210.1) without exon 8 which is probably co-amplified by reverse transcription PCR, the exact ratios of the mutant versus the normal splice product could not be determined. Unfortunately, it was not possible to design specific primers for the aberrant exon 8 skipped splice product. In a homozygous state, the *TANGO1* c.3621A>G mutation leads to exon 8 skipping in most splice products, whereas in the heterozygous parents the normal splice product is more abundant.

### The truncated TANGO1 protein does not localise to ER exit sites

To test the properties of the truncated TANGO1 protein, TANGO1 lacking exon 8 (Ex8-HA) was expressed in cultured HeLa cells, from which endogenous TANGO1 was knocked out (HeLaΔTANGO1) using CRISPR/Cas9 methodology (*Santos et al., 2015*). HeLaΔTANGO1 cells were transiently transfected with cDNA for either wild-type TANGO1-HA or Ex8-HA (*Figure 4*). 48h after transfection, cells were fixed, permeabilised and immunostained for HA and the ERES marker Sec16A. WT TANGO1-HA (green) expressed in distinct puncta, which colocalised with Sec16A (red). On the other hand, Ex8-HA (green) was distributed in a more diffused pattern throughout the cell and it did not localise to ERES (red) (*Figure 4A*). Transfected HeLaΔTANGO1 cells were probed with anti-HA and anti-Calreticulin (an ER-resident chaperone). WT TANGO1-HA (green) did not show any association with calreticulin (red), while Ex8-HA (green) was almost entirely colocalised with calreticulin (red) (*Figure 4B*). These data are consistent with our understanding of TANGO1 function, as its cytoplasmic domains are required to recruit TANGO1 to ERES. The Ex8 mutant lacks any cytoplasmic domains and consequently is distributed through the ER.

**Figure 4.**
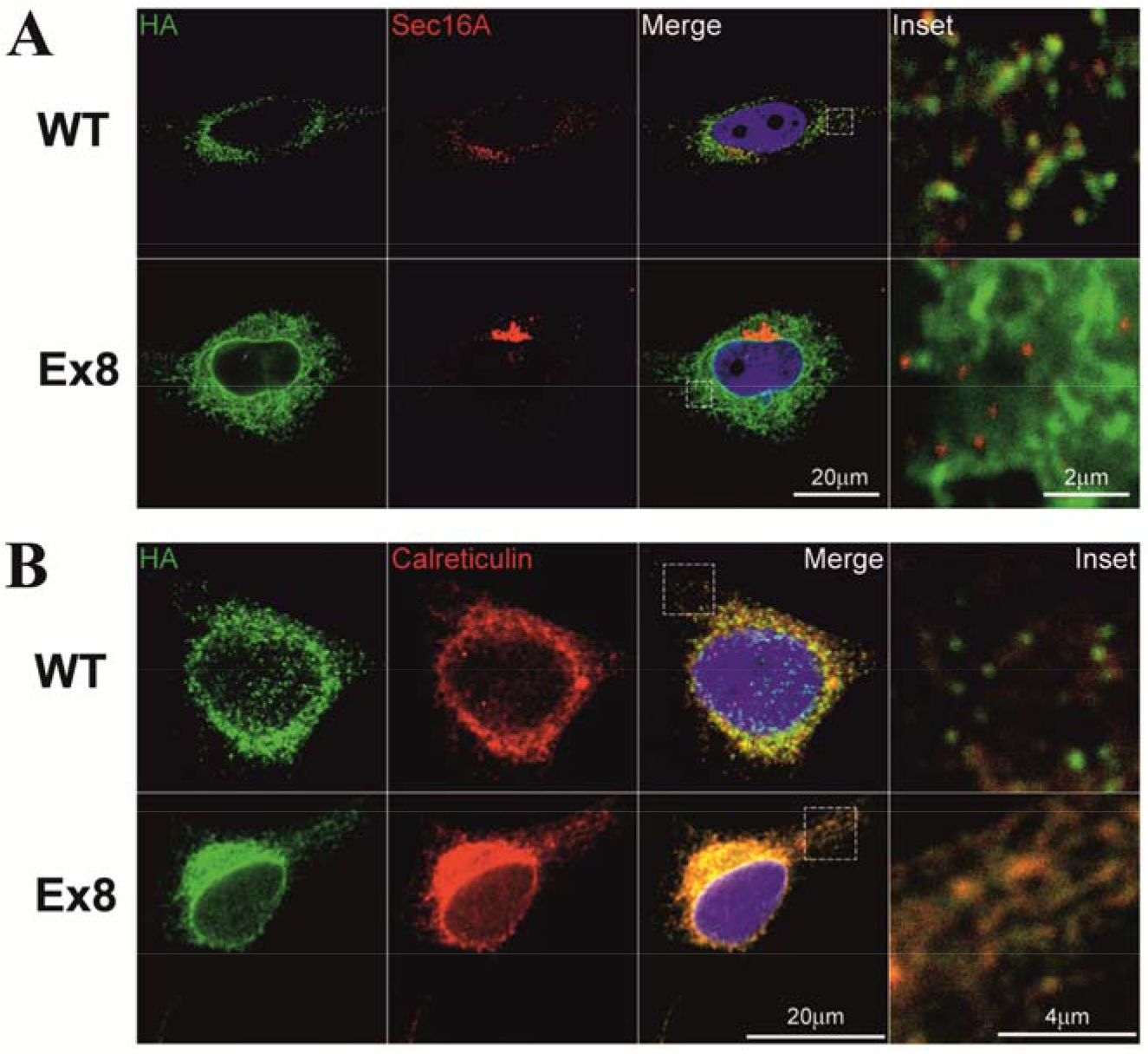
Ex8 mutant does not localise to ER exit sites. Immunofluorescence images of HeLa cells, lacking endogenous TANGO1, transiently transfected with WT TANGO1-HA (WT) or Ex8-HA (Ex8). (**A**) Cells were probed with anti-HA (green) and anti-Sec16A antibodies (red). Scale bar: 20 μm, inset 2 μm. (**B**) Cells were probed with anti-HA (green) and anti-Calreticulin (red). Scale bar 20 μm, inset 4 μm.

### Cells expressing the truncated TANGO1 show reduced levels of intracellular and secreted collagen I

All affected individuals showed the highest amounts of the exon 8 skipped splice product compared to the normal *TANGO1* splice product (*Figure 3*). However, it was not possible to determine how the relative abundance of the two splice products translated to protein levels in patient-derived samples. Therefore, possible effects of the overexpression of Ex8-HA on top of endogenous levels of normal *TANGO1* splice product at the cellular level were investigated using human osteosarcoma U2OS as a model system. U2OS cells produce and rapidly secrete collagen I, so they are the ideal system to monitor possible effects of Ex8-HA overproduction on collagen homeostasis. For this purpose, a stable U2OS cell line expressing Ex8-HA under a constitutive promoter was generated and compared with wild-type cells by microscopy. Cells expressing Ex8-HA showed weaker and more diffuse staining of collagen I compared with control cells (*Figure supplement 4A*). To confirm that this reduction was not due to unequal immunofluorescent staining, total RNA was isolated from the two cell populations and the relative amount of collagen I/GAPDH transcript was quantified by qRT-PCR analyses (*Figure supplement 4B*). This confirmed that collagen I expression is reduced in the Ex8-HA stable cell line.

The next step was to test whether the secretion of collagen I was also affected. Since different rates of collagen synthesis will affect the amount available for secretion in the two cell populations, a cycloheximide chase experiment was performed (*Figure 5*). By inhibiting protein synthesis, it was possible to monitor the rate of secretion of the available pool of collagen I present in the cells at time zero. By quantifying the relative amount of collagen I present in the cells and in the media at each time point, a drastic reduction in the rate of collagen I secretion from the EX8-HA stable cell line compared to control cells was observed. Importantly, this effect was not due to a general reduction of protein secretion since the small cargo antitrypsin was produced and secreted at a comparable rate in the two cell populations. Collectively, these results show that expression of the exon 8 skipped splice product even in the presence of full length TANGO1 affects collagen I homeostasis.

**Figure 5.**
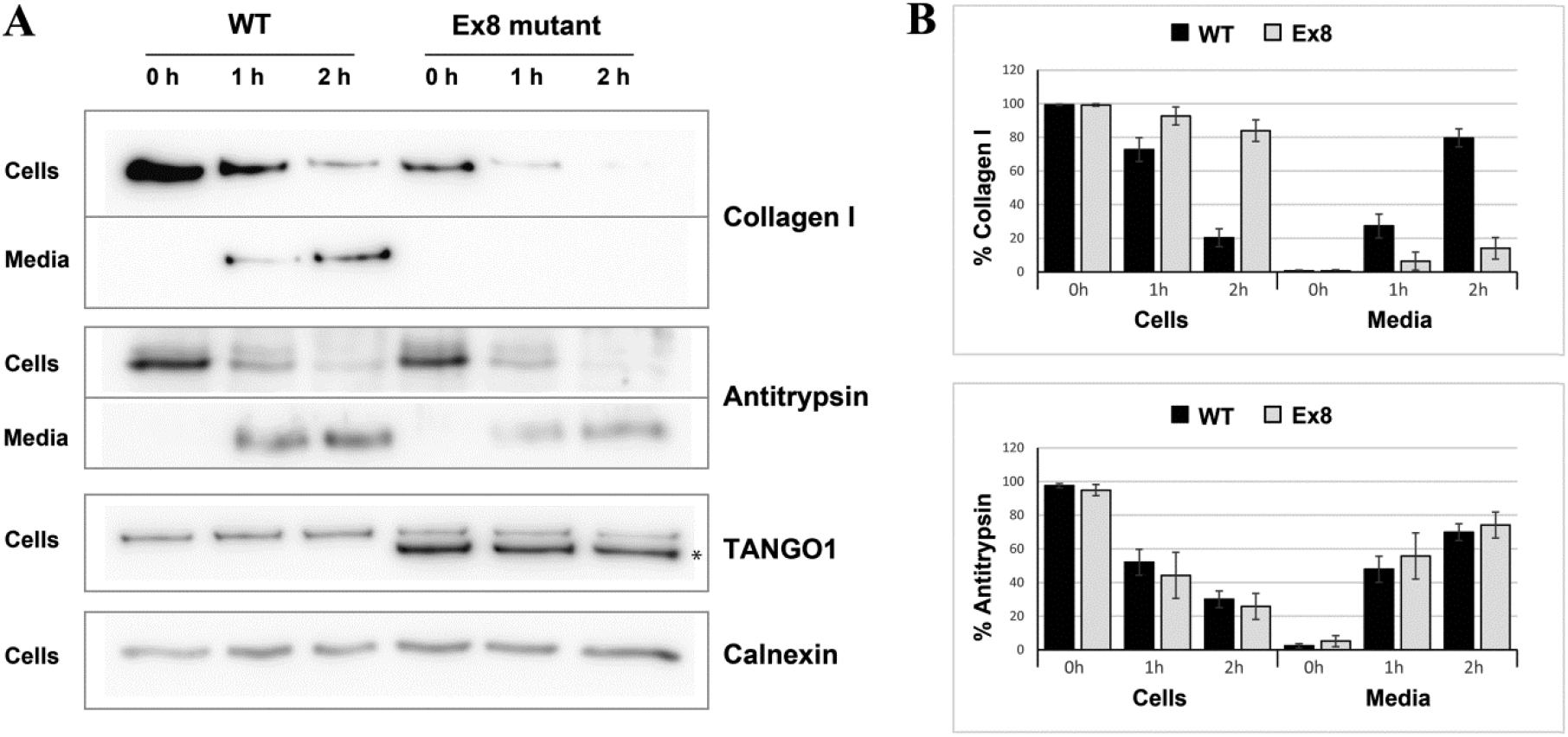
Ex8 mutant expression reduces collagen I secretion in U2OS cells. (**A**) Media of U2OS cells wild-type (WT) or stably expressing Ex8-HA mutant (Ex8) were replaced with OptiMEM media containing 0.25 mM ascorbic acid and 50 μM cycloheximide to block protein synthesis and follow collagen secretion. Cell extracts and media were collected at the indicated time points and analysed by SDS-PAGE followed by Western blotting with antibodies raised against Collagen I, TANGO1, Antitrypsin (small cargo) and Calnexin (loading control). (*) indicates Ex8-HA. (**B**) For each time point, the band intensities of collagen I (upper panel) or antitrypsin (lower panel) were measured for the cell extract and media samples and expressed as percentage of the total (cells plus media). Each graph represents the average quantification of four experiments and corresponding standard deviations.

## Discussion

Our study provides evidence that aberrant expression of a truncated TANGO1 protein and/or reduced levels of fully functional TANGO1 protein, or likely a combination of both causes a novel syndrome due to disturbances in cellular protein secretion. The heterozygous parents show exon 8 skipping but no detectable symptoms, whereas all four homozygous children are severely and similarly affected. This is consistent with a threshold model, where the disease only manifests when the ratio of truncated versus normal protein exceeds a critical level. *Tango1* knockout mice represent a full loss-of-function (LoF) situation and are defective for the secretion of numerous collagens, exhibiting short-limbed dwarfism, compromised chondrocyte maturation and bone mineralization, and other features (*Wilson et al., 2011*), resembling our patients’ phenotype. The human *TANGO1* locus lies within a homozygous interval shared among the affected children, which strengthens its role as the disease-causing gene in the investigated family. *TANGO1* does not seem to be haploinsufficient, since there are several heterozygous LoF mutation carriers listed in big population databases. However, no homozygous LoF mutation carriers have been reported so far. Thus, complete ablation of functional TANGO1 may cause embryonic lethality in humans.

The skipped exon 8 in full length TANGO1 corresponds to exon 3 in the isoform TANGO1-short. Its exclusion there also leads to a premature stop codon 27 bp downstream of exon 2. TANGO1-short has a similar structure to TANGO1, but lacks the lumenal portion within the cargo-binding SH3 domain. However, TANGO1-short has been shown to substitute the function of TANGO1 in collagen export, and vice versa (*Maeda et al., 2016*). It has been postulated that the cytoplasmic portion’s capacity to recruit ERGIC-53 membranes, Sec23/24 complexes, and cTAGE5, shared by both isoforms, is sufficient to export collagen at the ER (*Santos et al., 2015*; *Saito et al., 2009*). The lumenal SH3 domain may therefore rather play a role in modulating the efficiency or the quality (folding status) of collagens to be secreted (*Maeda et al., 2016*; *Raote et al., 2018*; *Saito et al., 2009*). Since the identified mutation in exon 8 does not only compromise the function of TANGO1, but most likely also of TANGO1-short, the short isoform cannot attenuate our patients’ phenotype.

The truncated TANGO1 protein is unable to access ERES and is distributed throughout the ER (*Figure 4*). Its expression in cells with endogenous full length TANGO1 displayed a negative effect on collagen secretion. This effect is specific to collagen, as the secretion of antitrypsin remains unaffected (*Figure 5*). It is conceivable that the truncated TANGO1 sequesters HSP47 away from ERES, which usually assists in collagen folding and export from the ER. It was recently shown that HSP47 may play a key role in modulating the unfolded protein response (UPR) by directly binding to the inositol-requiring enzyme 1 alpha (IRE1α) sensor (*Sepulveda et al., 2018*), thereby affecting the folding and the secretory capacity of the cell according to collagen production. An interesting speculation arising from our data would be that the HSP47/UPR axis has a bidirectional function - limiting collagen biogenesis when the secretory capacity is attenuated.

Type I collagens provide tensile strength to connective tissue and are abundant in bone, skin, dentin, cementum, tendons, and ligaments (*Desmukh et al., 2016*). Impaired collagen I secretion may underlie several key symptoms in our patients, in particular the tooth and skeletal abnormalities. However, the complex phenotype of the probands suggests that TANGO1 may also be involved in the secretion of further types of collagens, e.g. COL2A1, COL4A3, COL4A4, COL4A5, COL4A6, COL9A1, COL9A2 und COL9A3, COL11A1, and/or COL11A2, which are all important for inner ear function and associated with hereditary hearing loss (https://hereditaryhearingloss.org/). Defective secretion of other molecules, in particular hormones, may cause diabetes mellitus and pubertas praecox. Endocrinological examination revealed that the glucose intolerance of the affected children is due to reduced levels of secreted insulin. cTAGE5 is known to cooperate with TANGO1 in the mega cargo secretion pathway (*Saito et al, 2011)*, but has also been shown to play a pivotal role in ER to Golgi trafficking of small molecules like proinsulin (*Fan et al., 2017*). The knockout of *cTAGE5* in pancreatic β-cells resulted in defective islet structure, reduced insulin secretion, and severe glucose intolerance in mice (*Fan et al., 2017*). Additionally, a correlation between TANGO1 phosphorylation and proinsulin trafficking in mouse pancreatic β-cells has recently been discovered (*Kang et al., 2019*). In this light, it will be intriguing to investigate the role of TANGO1 in the insulin secretion pathway in future studies.

Collectively, the investigated family presents the first *TANGO1*-associated syndrome in humans, highlighting the role of fully functional TANGO1 in various disease pathways and bringing new potential target molecules of TANGO1 into focus.

## Materials and methods

### Whole exome sequencing

Exome capture was performed according to the Illumina Nextera Rapid Capture Enrichment library preparation (individuals II.1 and II.2) or the Illumina TruSeq Rapid Exome library preparation kit (individuals I.1, I.2, II.4, and II.5), using 50 ng of genomic DNA. Paired-end sequencing of the libraries was performed with a NextSeq500 sequencer and the v2 reagent kit (Illumina, San Diego, California, USA). Sequences were mapped to the human genome reference (NCBI build37/hg19 version) using the Burrows-Wheeler Aligner. Aligned reads ranged between 82,649,383 and 102,537,469. The mean coverage was ≥52 with 90.3% of the exome being covered at least 10x. A total of 237,330-297,312 variants per sample were called and analyzed using GensearchNGS software (PhenoSystems SA, Braine le Chateau, Belgium). Variants with a coverage of ≤20, a Phred-scaled quality of ≤15, a frequency of ≤20, and a MAF of ≥1% were neglected. Two control samples from healthy individuals were used for filtering out platform artefacts. Alamut Visual (Interactive Biosoftware, Rouen, France) software including prediction tools like SIFT, MutationTaster, and PolyPhen-2 was used for variant prioritization. Potential effects of a variant on pre-mRNA splicing were evaluated by SpliceSiteFinder-like, MaxEntScan, NNSPLICE, GeneSplicer, Human Splicing Finder, ESEfinder, RESCUE-ESE, and EX-SKIP. Population databases like ExAC, gnomAD, and GME revealed whether a variant has been previously found. Protein expression, structure, and functional aspects were investigated with UniProt and The Human Protein Atlas. Information on mouse models was retrieved from the MGI database.

### Sanger sequencing

*TANGO1* exon 8 was amplified by a touchdown PCR program using primers in the flanking introns (forward 5’-TCAGACCACAACATATCACTACTGG-3’; reverse 5’-TACTCTATCATACAACCTGGCAACC-3’). A clean-up step with ExoSAP-IT (Applied Biosystems, Foster City, California, USA) was followed by the sequencing reaction using the BigDye Terminator Cycle Sequencing Kit v1.1 (Applied Biosystems). Sequencing was conducted on a 3130XL capillary sequencer (Applied Biosystems) and data analysis was performed with Gensearch (PhenoSystems SA).

### Microarray analyses

DNA from two affected children (II.1 and II.2) was genotyped using the Infinium Global Screening Array-24 v1.0 BeadChip (Illumina) by Life & Brain (Bonn, Germany). Shared homozygous intervals were identified with HomozygosityMapper (*Seelow et al., 2009*).

Array CGH was performed using the CGX DNA labeling kit (PerkinElmer, Waltham, Massachusetts, USA) and the CGX-HD array (PerkinElmer) that covers clinically relevant regions with 180,000 oligonucleotide marker. A male genomic DNA sample served as a reference. The hybridized array was scanned with the NimbleGen MS 200 Microarray Scanner (Roche, Basel, Switzerland). Data analysis was conducted with CytoGenomics 2.5 (Agilent Technologies, Santa Clara, California, USA) and Genoglyphix 3.0 (PerkinElmer) software using annotations from GRCh37/hg19.

### Minigene assay

To investigate possible effects of the *TANGO1* mutation on pre-mRNA splicing, the homozygous individual II.1 was compared to a normal control sample in a minigene assay. The plasmid pSPL3b-cam vector (*Burn et al., 1995*) is endowed with a chloramphenicol resistance, an SV40 promoter, SD6 and SA2 primer sequences, as well as a multiple cloning site including recognition sites for *XhoI* and *BamHI* (*Figure supplement 2A*). A 521 bp amplicon including *TANGO1* exon 8 and ∼250 bp flanking intronic sequences was generated from genomic DNAs of the patient and a control, using primers with recognition sites for *XhoI* and *BamHI* at their 5’ ends (fwd 5’-AATTCTCGAGTATCTTTAGCTGTGCAAAGT-3’; rev 5’-ATTGGATCCAAGGTCAATCTGCCCCAAAT-3’) and the Q5 High-Fidelity DNA Polymerase (New England Biolabs, Ipswich, Massachusetts, USA). The PCR products were purified with the GenElute PCR Clean-Up kit (Sigma-Aldrich, St. Louis, Missouri, USA), digested by *XhoI* and *BamHI* in CutSmart Buffer (New England Biolabs), again purified, and finally ligated into the linearized vector using T4 DNA Ligase and T4 DNA Ligase Reaction Buffer (New England Biolabs).

Vector constructs were transformed into DH5α bacteria by heat shock for 90 sec at 42°C and then plated onto LB/agar/chloramphenicol Petri dishes. Following overnight incubation at 37°, a colony screen was performed using SD6 (fwd 5’-TCTGAGTCACCTGGACAACC-3’) and the *TANGO1* exon 8 reverse primer (see above). Positive clones (with insert) were cultured overnight and the vector constructs extracted using the GenElute Plasmid Miniprep kit (Sigma-Aldrich). Sequencing was performed with 100 fmol of vector constructs, SD6 forward and *TANGO1* exon 8 reverse primers. Three vector constructs were selected for splicing experiments, one with the wild-type *TANGO1* exon 8 and flanking sequences (WT vector) and another one with the mutated *TANGO1* exon 8 (Mnt vector). The pSPL3b-cam vector without insert (CTRL) served as control.

Aliquots of 4×10^15^ HEK293T cells were plated into 6-well-plates and transfected with vector constructs (WT, Mnt, or CTRL) using the FuGENE HD Transfection Reagent (Roche). After 24 h of incubation at 37°C, RNA was isolated with the miRNeasy Mini kit (Qiagen, Venlo, Netherlands). cDNA was synthesized using the High Capacity RNA-to-cDNA kit (Applied Biosystems), amplified by a touchdown PCR program using SD6 forward and SA2 reverse (5’-ATCTCAGTGGTATTTGTGAGC-3’) primers, purified with ExoSAP-IT (Applied Biosystems) and finally sequenced. The resulting splice products were compared by gel electrophoresis and cDNA sequencing. Since the Mnt vector produced two separate *TANGO1* splice products, individual cDNA molecules were cloned using the TA Cloning Kit and the Dual Promoter (pCRII) protocol (Invitrogen, Carlsbad, California, USA).

### RNA isolation from whole blood samples, cDNA synthesis and sequencing

Peripheral blood samples were collected in PAXgene Blood RNA tubes and RNA was isolated using the PAXgene Blood RNA Kit (PreAnalytiX, Hombrechtikon, Switzerland). Reverse transcription was performed with the High Capacity RNA-to-cDNA Kit (Applied Biosystems). cDNA was amplified by a touchdown PCR program using primers located in *TANGO1* exon 6 and 11, respectively (fwd 5’-ACACTCCTATGGATGCTATTGATGC-3’; rev 5’-CTCTCTCAGATTCTAGCATAACACG-3’). After a clean-up step with ExoSAP-IT (Applied Biosystems), cDNA sequencing was performed with the BigDye Terminator Cycle Sequencing Kit v1.1 (Applied Biosystems) and a 3130XL capillary sequencer (Applied Biosystems). Data analysis was performed with Gensearch (PhenoSystems SA) and CodonCode Aligner (CodonCode Corporation, Centerville, Massachusetts, USA). Multiple PCR products were separated by gel electrophoresis. cDNA of individual cut out bands was isolated with the QIAquick Gel Extraction Kit (Qiagen) and 1-3 µl of gel extracts were used for sequencing.

### Morpholino assays

Effects of different *vivo* morpholinos on *TANGO1* pre-mRNA splicing were tested on HeLa cells with a different genetic background. 2×10^5^ HeLa cells in 2 ml DMEM (Sigma-Aldrich) each were plated into the required number of wells of a 6-well-plate and incubated at 37°C for 24 h. Depending on the research question, cells were then transfected with the *TANGO1* WT or Mnt vector, or used without vector transfection for morpholino treatment. For transfection, 2 µg vector were suspended in up to 95 µl DMEM and mixed with 6 µl FuGENE HD Transfection Reagent (Roche). The mixture was incubated at room temperature for 15 min and then slowly added to the cells.

The customized TGO morpholino targets the entire *TANGO1* exon 8. A standard control morpholino (GeneTools, Philomath, Oregon, USA) was used to exclude unspecific effects. The *vivo* morpholinos were added at a final concentration of 5 or 7 µM. After 24 h incubation, the medium was removed, cells were washed with 2 ml 1xPBS, detached with 0.5 ml Trypsin-EDTA Solution (Sigma-Aldrich) and then transferred into 2 ml tubes with 1x PBS. Samples were centrifuged at 3000 g for 5 min at 4°C. Supernatant was removed and RNA isolated using the miRNeasy Mini Kit (Qiagen). cDNA synthesis and sequencing were performed as described above, using vector-specific primers (SD6 and SA2). This ensured that only vector-derived splice products were analysed.

### Quantitative real-time PCR

qRT-PCRs were performed to obtain a relative ratio of either the exon 8 skipped *TANGO1* or the normal splice product for homozygous and heterozygous mutation carriers, compared to a reference sample (without mutation). Two housekeeping genes, *HPRT1* (fwd 5’-TGACACTGGCAAAACAATGCA-3’; rev 5’-GGTCCTTTTCACCAGCAAGCT-3’) and *IPO8* (fwd 5’-CGAGCTAGATCTTGCTGGGT-3’; rev 5’-CGCTAATTCAACGGCATTTCTT-3’) served as endogenous controls. Assay 1 (fwd 5’-GACTGCCATGGAAACCTGTATT-3’; rev 5’-TCCGTGAAACAAGGACAGTTCT-3’) exclusively amplified the mutant splice product where exon 7 is followed by exon 9; assay 2 (fwd 5’-GACTGCCATGGAAACCTGTATT-3’; rev 5’-CCTTCACAACAAGGACAGTTCT-3’) the normal splice products where exon 7 is followed by exon 8. The PCR reaction consisted of 4 µl (10 ng) cDNA, 1 µl (2.5 pmol) primer pair, 2 µl 5x HOT FIREPol EvaGreen qPCR Mix Plus, and 3 µl water. All samples were run in technical triplicates. Cycling conditions on a ViiA 7 Real-Time PCR System (Applied Biosystems) were as follows: 95°C for 15 min, 40 cycles of 95°C for 15 sec, 60°C for 20 sec, and 72°C for 20 sec. The melt curve was obtained from 60°C to 95°C and indicated no secondary amplicons.

In addition, qRT-PCRs were conducted to measure the relative expression of full length TANGO1, truncated TANGO1, and Collagen I. For this purpose, U2OS cells WT or stabling expressing Ex8-HA were lysed and total RNA extracted with the RNeasy extraction kit (Qiagen). cDNA was synthesized with Superscript III (Invitrogen). Primers for collagen I alpha-chain (fwd 5’-GTGGTCAGGCTGGTGTGATG-3’; rev 5’-CAGGGAGACCCTGGAATCCG-3’), TANGO1 lumenal portion (fwd 5’-TGGAAGTGTTGGACGCACTTTT-3’; rev 5’-TCAGGTTCAGGTTCCCTTTCCT-3’) or cytosolic portion (fwd 5’-CTCAGCTCTGCGGACCTTTT3’; rev 5’-GTGAACAGTCCTGGCTAGTGC-3’) were designed using Primer-BLAST (NCBI) (Ye et al., 2012) with the annealing temperature to 60°C. To determine expression levels of collagen I and the two forms of TANGO1, qRT-PCR was performed with Light Cycler 480 SYBR Green I Master (Roche) according to manufacturer’s instructions.

### Cell culture and transfection

U2OS and RDEB/C7 cells were grown at 37°C with 5% CO_2_ in complete DMEM with 10% FBS. For lentiviral infection of Ex8-HA into U2OS and RDEB/C7 cells, lentiviral particles were produced by cotransfecting HEK293 cells with pHRSIN/Ex8-HA plasmid and a third-generation packaging vector pool using TransIT-293 (Mirus Bio, Madison, Wisconsin, USA). 72 h after transfection, the viral supernatant was harvested, filtered, and directly added to U2OS and RDEB/C7 cells. Infected cells were selected using 500 µg/ml hygromycinB (Invitrogen). The TANGO1-knockout HeLa cell line was described previously (*Santos et al., 2015*).

### Collagen-secretion assays

The media of U2OS cells was replaced with Optimem medium (Thermo Fisher Scientific, Waltham, Massachusetts, USA) containing 0.25 mM ascorbic acid and 50 µM cycloheximide (Sigma-Aldrich) for up to 2 h to allow for collagen secretion. The media were collected at 0, 1, and 2 h time points, centrifuged at low speed to remove any cells or cellular debris, and the supernatants were denatured at 65°C for 10 min with Laemmli SDS sample buffer. For cell extracts, cells were washed with PBS, lysed in buffer A (50 mM Tris-Cl, pH7.4, 150 mM NaCl, 1 mM EDTA, 1% Triton X-100) plus proteases inhibitors (Roche), and centrifuged at 14,000 rpm for 15 min at 4°C. The supernatants were denatured at 65°C for 10 min with Laemmli SDS sample buffer. Media and cell lysate were subjected to SDS-PAGE (6% or 8% acrylamide) and Western blotting with antibodies raised against collagen I, TANGO1, calnexin, and antitrypsin. Band intensities were measured using QuantityOne (Bio-Rad, Hercules, California, USA), and four repetitions of the experiment were used to plot the graph.

### Immunofluorescence staining

Cells grown on coverslips were fixed with cold methanol for 10 min at −20°C and incubated with blocking reagent (Roche) for 30 min at RT. Primary antibodies were diluted in blocking reagent and incubated overnight at 4°C. Secondary antibodies conjugated with Alexa Fluor 488 or 594 (Invitrogen) were diluted in blocking reagent and incubated for 1 h at room temperature. Images were taken with a TCS SP8 or TCS SPE confocal microscope (Leica Microsystems, Wetzlar, Germany) with a 63× objective. Images processing was performed with ImageJ.

### Antibodies

Antibodies used in Western blotting and immunofluorescence microscopy were as follows: TANGO1 (Sigma-Aldrich); calreticulin (Novus Biologicals, Centennial, Colorado, USA); antitrypsin Ab-1 (NeoMarkers, Fremont, California, USA); collagen I (Abcam, Cambridge, United Kingdom); Sec16A (Sigma-Aldrich); rat hemagglutinin (Roche) or mouse hemagglutinin (Santa Cruz Biotechnology, Dallas, Texas, USA).

## Acknowledgement

We would like to thank the family for their participation.

## Author contributions

C.L. performed exome sequencing and data analysis, *in vivo* and *in vitro* splice assays, qRT-PCR analyses, and morpholino experiments. O.F. and I.R. conducted functional studies on effects of the *TANGO1* mutation on protein localisation and collagen I homeostasis. D.L. supervised the morpholino experiments and the qRT-PCR analyses. E.M.K. supervised the *in vitro* splice assay. I.N. performed aCGH analysis. B.V. contributed conceptual input. P.D.C. and R.C. provided clinical data and patient samples. V.M. and T.H. designed the study. C.L., V.M. and T.H. wrote the manuscript and all co-authors reviewed the manuscript.

---

**Figure supplement 1.**
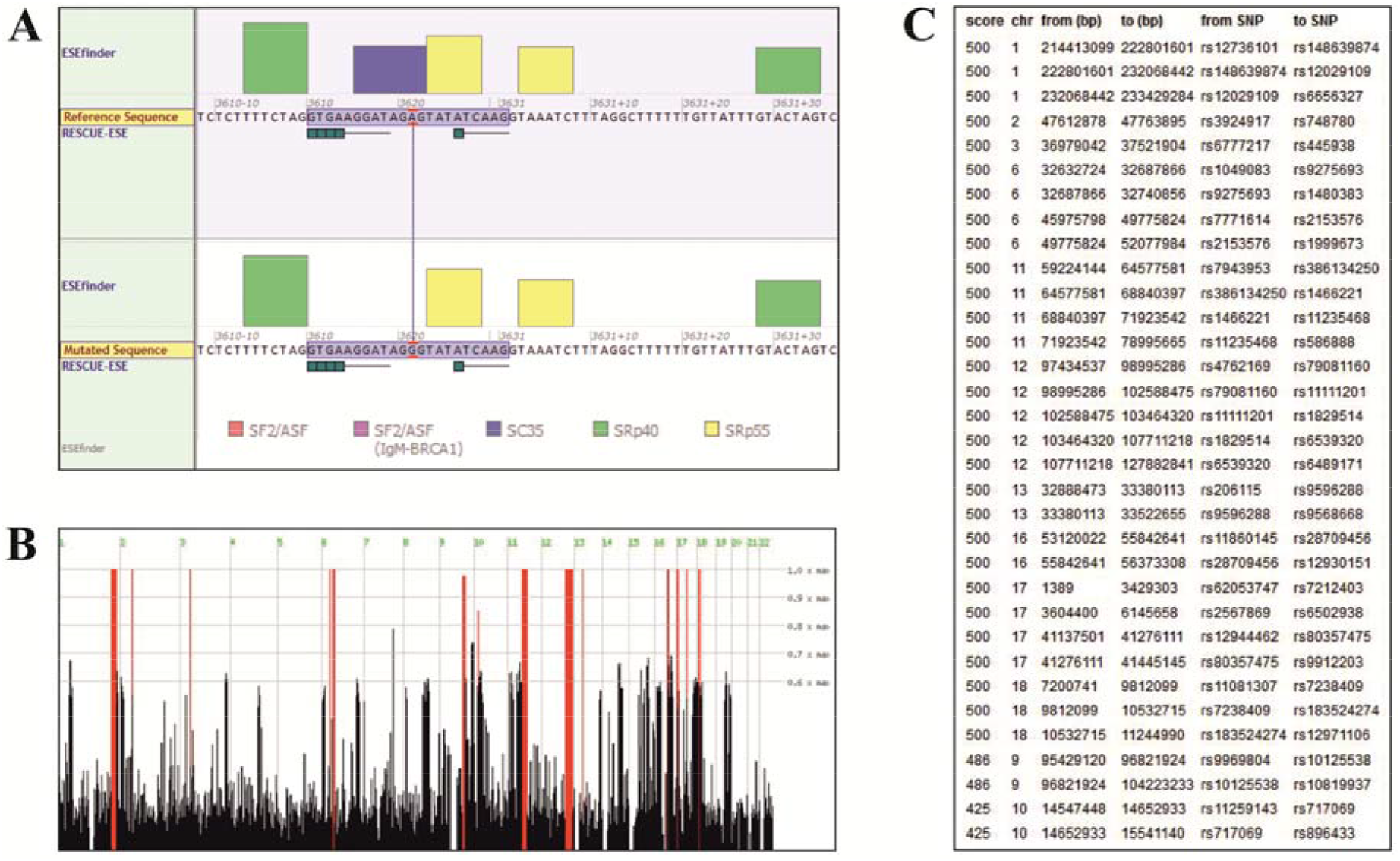
Sequence analysis. (**A**) Splicing of *TANGO1* exon 8, as predicted by Alamut Visual. The window shows the binding sites of various SR proteins (color code at the bottom) required for correct splicing. The height of the boxes reflects the probability of binding. The upper diagram shows the WT, the lower half the mutated sequence. The mutation is predicted to disrupt the consensus sequence for the SR protein SC35. (**B**) Homozygous intervals (red bars) shared by patients II.1 and II.2 of the investigated family. The identified *TANGO1* mutation lies within a ∼19 Mb homozygous interval on chromosome 1. (**C**) Genomic localisation (GRCh37/hg19) of homozygous intervals, detected by HomozygosityMapper.

**Figure supplement 2.**
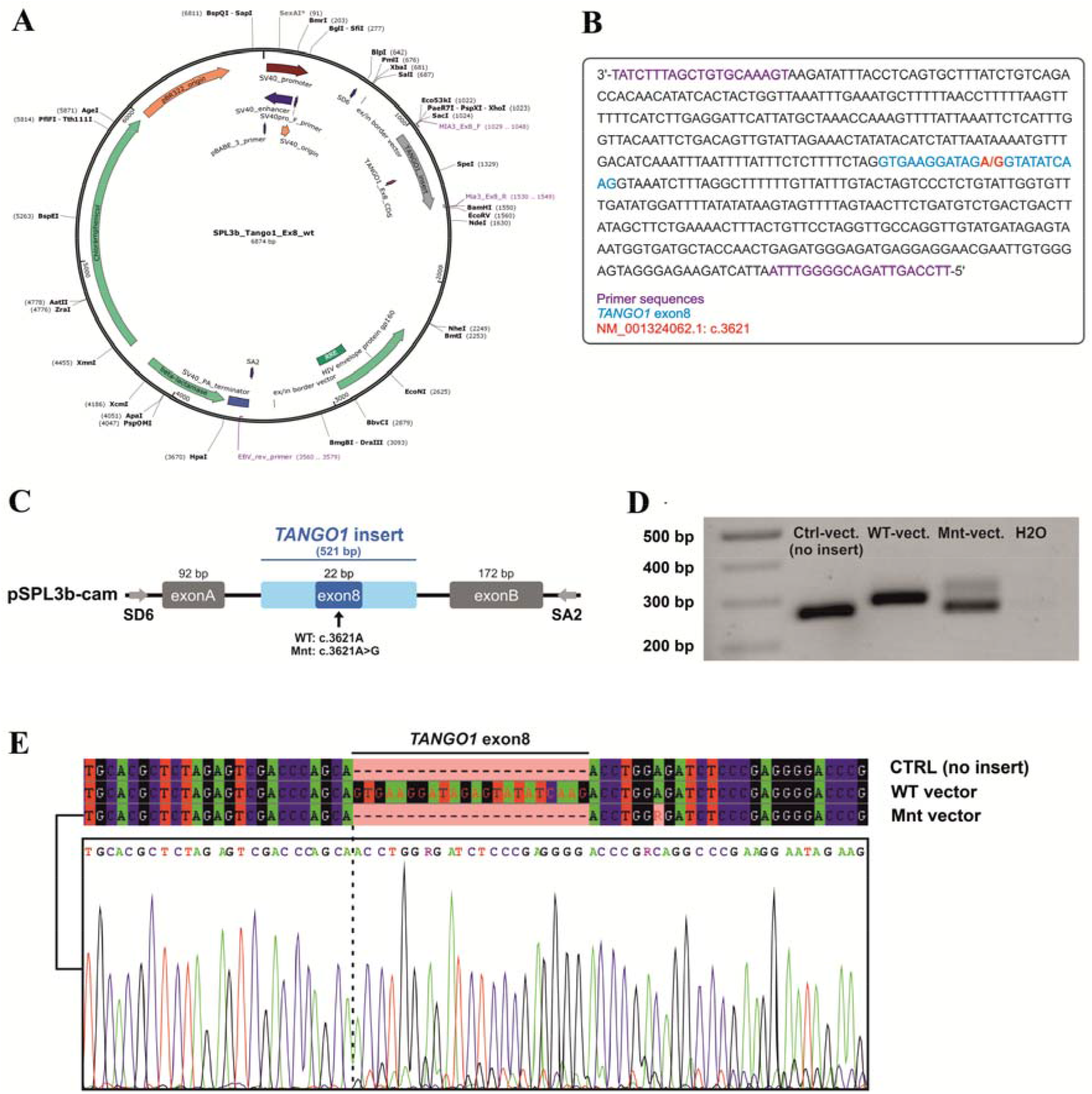
Minigene assay. (**A**) Map of the pSPL3b-cam vector. (**B**) Sequence of the 521 bp amplicon (*TANGO1* exon 8 and ∼250 bp flanking intronic sequences) inserted into pSPL3b-cam. (**C**) Vector constructs for the minigene assay. An amplicon containing either wild-type (WT vector) or mutated exon 8 (Mnt vector) was inserted between the vector-specific exons A and B into pSPL3b-cam. (**D**) Gel electrophoresis of cDNA PCR products from HEK293T cells transfected with either WT, Mnt, or CTRL vector (without insert). Transfection with the CTRL and WT vector resulted in cDNA PCR products of 264 bp and 286 bp, respectively. Cells transfected with the Mnt vector produced two splice products. (**E**) Sanger cDNA sequencing of the CTRL-, WT-, and Mnt-vector splice products. The Mnt vector yielded two separate *TANGO1* splice products, the majority of which did not contain exon 8, indicative of exon skipping and a small portion of correctly spliced products. Following TA cloning of individual cDNA molecules, 15 of 22 (68%) splice products lacked and 7 (32%) contained exon 8.

**Figure supplement 3.**
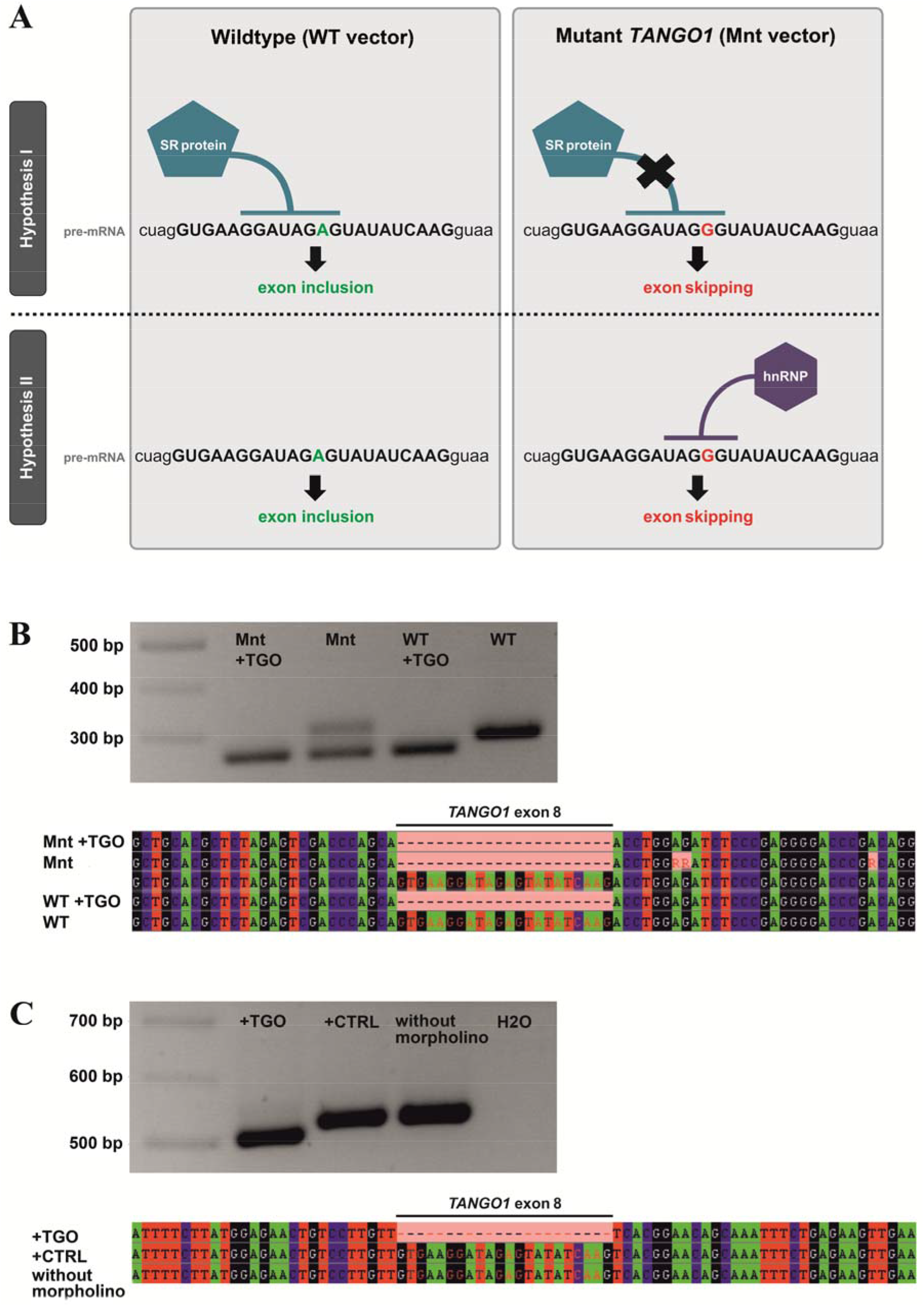
(**A**) Possible mechanisms underlying *TANGO1* exon 8 skipping. Exon skipping during pre-mRNA splicing could be due to disruption of an ESE motif, which prevents the human SR protein SC35 from binding (hypothesis I). On the other hand, the mutation creates a sequence motif (UAGGGU) that is recognised by hnRNP A1, which may repress splicing (hypothesis II). Most morpholinos alter splicing by sterically blocking the snRNP binding sites utilised by the spliceosome, which are usually located in the intron near the splice donor/acceptor junctions. The TGO morpholino used here targets the entire *TANGO1* exon 8 but not enough intronic sequences to block snRNP binding sites. It rather obstructs the consensus sequence for SC35 binding. If disruption of a splice enhancer motif is the underlying mechanism, TGO treatment of WT-transfected cells should also prevent SC35 binding and thus induce exon 8 skipping. If a splice inhibitor is recruited by the mutated sequence, TGO treatment of WT-transfected cells should not affect *TANGO1* exon 8 splicing. In cells expressing the mutation from a transfected vector, addition of the TGO morpholino would prevent the binding of hnRNP A1 and induce exon 8 skipping. (**B**) Effects of the *TANGO1* exon 8 morpholino (TGO) on HeLa cells transfected with mutated *TANGO1* exon 8 (Mnt vector) or wild-type (WT vector). To discriminate between vector-derived and endogenous splice products, vector-specific primers were used for cDNA sequencing. The vector-derived splice products of TGO treated HeLa cells transfected with either mutated (Mnt) or wild-type (WT) *TANGO1* both showed exon skipping. HeLa cells transfected with the Mnt vector but not with TGO were endowed with two splice products, one lacking the entire *TANGO1* exon 8 and one presenting the normal cDNA. Cells transfected with the WT vector but not with TGO demonstrated only the normal *TANGO1* splice product. (**C**) To proof that the exon skipping effect is not caused by morpholino treatment alone, HeLa cells were either treated with TGO or a standard control morpholino (CTRL). As expected, cDNA from TGO-treated cells lacked *TANGO1* exon 8, whereas cDNAs from CTRL-treated or untreated cells included exon 8. Collectively, these results suggest that the identified *TANGO1* mutation leads to the disruption of an exon splice enhancer, which prevents SC35 binding.

**Figure supplement 4.**
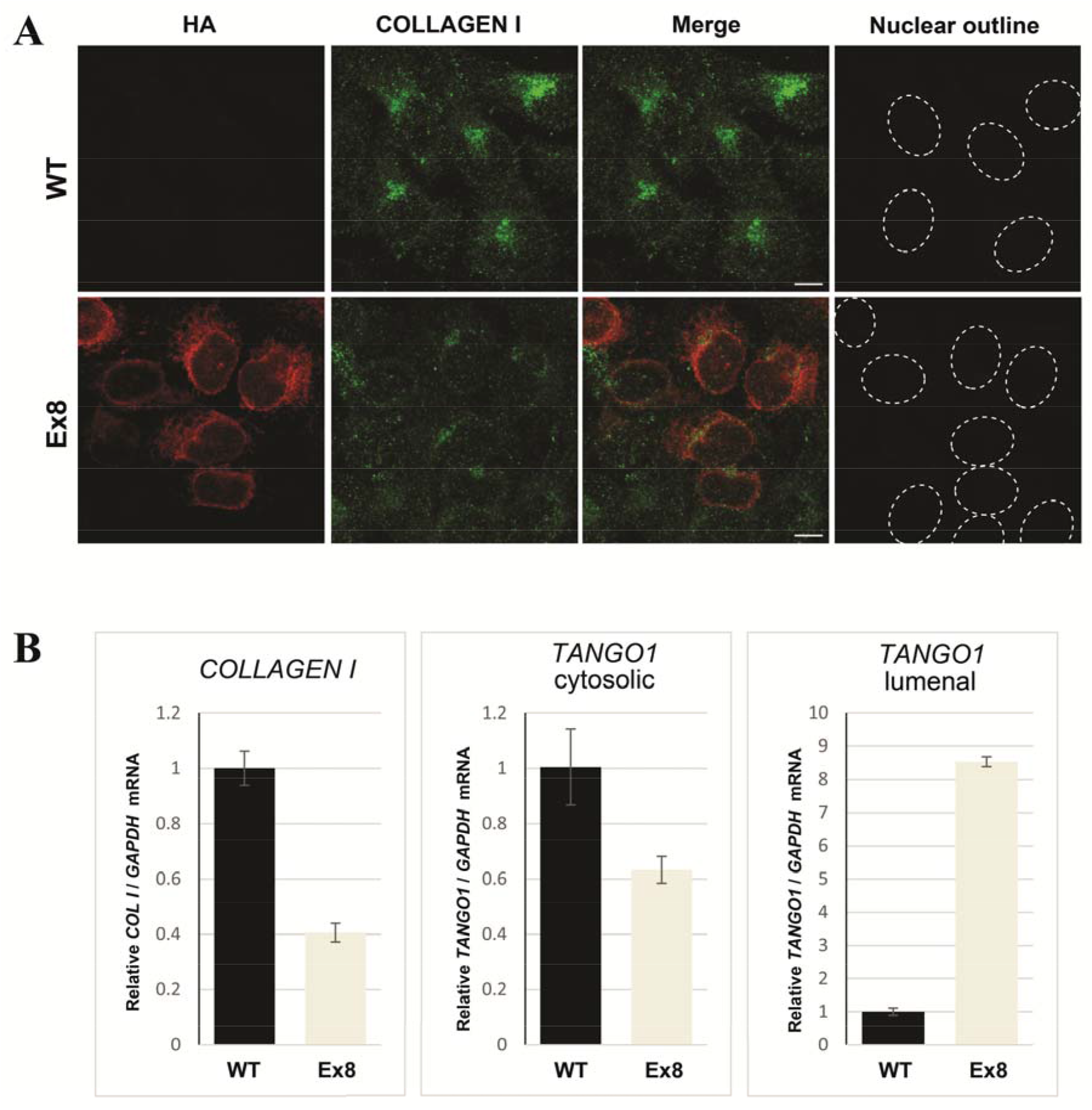
TANGO1 exon 8 mutant (Ex8) expression reduces collagen I expression in U2OS cells. (**A**) Immunofluorescence Z-stack projections of control wild-type or Ex8-HA expressing U2OS cells, probed with anti-HA antibody (red) and anti-Collagen I antibody (green). Nuclear borders were traced from DIC images. Scale bar = 10 µm. (**B**) RNA levels from control wild-type or Ex8-HA expressing U2OS cells normalised by *GAPDH* values. Primers were designed to amplify two different portions of *TANGO1* mRNA corresponding to the cytosolic portion (to quantify mRNA of endogenous protein only) or to the lumenal portion of TANGO1 protein (to quantify mRNA of endogenous *TANGO1* and overexpressed Ex8-HA). Collagen I primers were specific for alpha-chain mRNA.

## References

Bard F, Casano L, Mallabiabarrena A, Wallace E, Saito K, Kitayama H, Guizzunti G, Hu Y, Wendler F, Dasgupta R, Perrimon N, Malhotra V. 2006. Functional genomics reveals genes involved in protein secretion and Golgi organization. Nature 439:604–607.

Bosserhoff AK, Moser M, Schölmerich J, Buettner R, Hellerbrand C. 2003. Specific expression and regulation of the new melanoma inhibitory activity-related gene MIA2 in hepatocytes. Journal of Biological Chemistry 278:15225–15231.

Burn TC, Connors TD, Klinger KW, Landes GM. 1995. Increased exon-trapping efficiency through modifications to the pSPL3 splicing vector. Gene 161:183–187.

Cauwels RG, De Coster PJ, Mortier GR, Marks LA, Martens LC. 2005. Dentinogenesis imperfecta associated with short stature, hearing loss and mental retardation: a new syndrome with autosomal recessive inheritance? Journal of Oral Pathology & Medicine 34:444–446.

Deshmukh SN, Dive AM, Moharil R, Munde P. 2016. Enigmatic insight into collagen. Journal of Oral and Maxillofacial Pathology 20:276–283.

Fan J, Wang Y, Liu L, Zhang H, Zhang F, Shi L, Yu M, Gao F, Xu Z. 2017. *cTAGE5* deletion in pancreatic β cells impairs proinsulin trafficking and insulin biogenesis in mice. Journal of Cell Biology 216:4153–4164.

Ishikawa Y, Ito S, Nagata K, Sakai LY, Bächinger HP. 2016. Intracellular mechanisms of molecular recognition and sorting for transport of large extracellular matrix molecules. PNAS 113:E6036–E6044.

Kang T, Boland BB, Alarcon C, Grimsby JS, Rhodes CJ, Larsen MR. 2019. Proteomic analysis of restored insulin production and trafficking in obese diabetic mouse pancreatic islets following euglycemia. Journal of Proteome Research Epub ahead of print: doi 10.1021/acs.jproteome.9b00160.

Maeda M, Saito K, Katada T. 2016. Distinct isoform-specific complexes of TANGO1 cooperatively facilitate collagen secretion from the endoplasmic reticulum. Molecular Biology of the Cell 27:2688–2696.

Malhotra V, Erlmann P. 2011. Protein export at the ER: loading big collagens into COPII carriers. EMBO Journal 30:3475–3480.

Malhotra V, Erlmann P. 2015. The pathway of collagen secretion. Annual Review of Cell and Developmental Biology 31:109–124.

Malhotra V, Erlmann P, Nogueira C. 2015. Procollagen export from the endoplasmic reticulum. Biochemical Society Transactions 43:104–107.

Miller EA, Schekman R. 2013. COPII - a flexible vesicle formation system. Current Opinion in Cell Biology 25:420–427.

Raote I, Malhotra V. 2019. Protein transport by vesicles and tunnels. Journal of Cell Biology 218:737–739.

Raote I, Ortega Bellido M, Pirozzi M, Zhang C, Melville D, Parashuraman S, Zimmermann T, Malhotra V. 2017. TANGO1 assembles into rings around COPII coats at ER exit sites. Journal of Cell Biology 216:901–909.

Raote I, Ortega-Bellido M, Santos AJ, Foresti O, Zhang C, Garcia-Parajo MF, Campelo F, Malhotra V. 2018. TANGO1 builds a machine for collagen export by recruiting and spatially organizing COPII, tethers and membranes. Elife 7:e32723.

Saito K, Chen M, Bard F, Chen S, Zhou H, Woodley D, Polischuk R, Schekman R, Malhotra V. 2009. TANGO1 facilitates cargo loading at endoplasmic reticulum exit sites. Cell 136:891–902.

Saito K, Yamashiro K, Ichikawa Y, Erlmann P, Kontani K, Malhotra V, Katada T. 2011. cTAGE5 mediates collagen secretion through interaction with TANGO1 at endoplasmic reticulum exit sites. Molecular Biology of the Cell 22:2301–2308.

Santos AJ, Raote I, Scarpa M, Brouwers N, Malhotra V. 2015. TANGO1 recruits ERGIC membranes to the endoplasmic reticulum for procollagen export. Elife 4:e10982.

Santos AJ, Nogueira C, Ortega-Bellido M, Malhotra V. 2016. TANGO1 and Mia2/cTAGE5 (TALI) cooperate to export bulky pre-chylomicrons/VLDLs from the endoplasmic reticulum. Journal of Cell Biology 213:343–354.

Seelow D, Schuelke M, Hildebrandt F, Nürnberg P. 2009. HomozygosityMapper - an interactive approach to homozygosity mapping. Nucleic Acids Research 37:W593–599.

Sepulveda D, Rojas-Rivera D, Rodríguez DA, Groenendyk J, Köhler A, Lebeaupin C, Ito S, Urra H, Carreras-Sureda A, Hazari Y, Vasseur-Cognet M, Ali MMU, Chevet E, Campos G, Godoy P, Vaisar T, Bailly-Maitre B, Nagata K, Michalak M, Sierralta J, Hetz C. 2018. Interactome screening identifies the ER lumenal chaperone Hsp47 as a regulator of the unfolded protein response transducer IRE1α. Molecular Cell 69:238–252.e7.

Wilson DG, Phamluong K, Li L, Sun M, Cao TC, Liu PS, Modrusan Z, Sandoval WN, Rangell L, Carano RA, Peterson AS, Solloway MJ. 2011. Global defects in collagen secretion in a Mia3/TANGO1 knockout mouse. Journal of Cell Biology 193:935–951.

Ye J, Coulouris G, Zaretskaya I, Cutcutache I, Rozen S, Madden TL. 2012. Primer-BLAST: a tool to design target-specific primers for polymerase chain reaction. BMC Bioinformatics 13:134.

